# Impact of elevated IOP on lamina cribrosa oxygenation; A combined experimental-computational study on monkeys

**DOI:** 10.1101/2024.09.05.609208

**Authors:** Yuankai Lu, Yi Hua, Bingrui Wang, Fuqiang Zhong, Andrew Theophanous, Shaharoz Tahir, Po-Yi Lee, Ian A. Sigal

## Abstract

**Purpose:** Our goal is to evaluate how lamina cribrosa (LC) oxygenation is affected by the tissue distortions resulting from elevated IOP.

**Design:** Experimental study on monkeys

**Subjects:** Four healthy monkey eyes with OCT scans with IOP of 10 to 50 mmHg, and then with histological sections of LC.

**Methods:** Since in-vivo LC oxygenation measurement is not yet possible, we used 3D eye-specific numerical models of the LC vasculature which we subjected to experimentally-derived tissue deformations. We reconstructed 3D models of the LC vessel networks of 4 healthy monkey eyes from histological sections. We also obtained in-vivo IOP-induced tissue deformations from a healthy monkey using OCT images and digital volume correlation analysis techniques. The extent that LC vessels distort under a given OCT-derived tissue strain remains unknown. We therefore evaluated two biomechanics-based mapping techniques: cross-sectional and isotropic. The hemodynamics and oxygenations of the four vessel networks were simulated for deformations at several IOPs up to 60mmHg. The results were used to determine the effects of IOP on LC oxygen supply, assorting the extent of tissue mild and severe hypoxia.

**Main Outcome Measures:** IOP-induced deformation, vasculature structure, blood supply, and oxygen supply for LC region

**Result:** IOP-induced deformations reduced LC oxygenation significantly. More than 20% of LC tissue suffered from mild hypoxia when IOP reached 30 mmHg. Extreme IOP(>50mmHg) led to large severe hypoxia regions (>30%) in the isotropic mapping cases.

**Conclusion:** Our models predicted that moderately elevated IOP can lead to mild hypoxia in a substantial part of the LC, which, if sustained chronically, may contribute to neural tissue damage. For extreme IOP elevations, severe hypoxia was predicted, which would potentially cause more immediate damage. Our findings suggest that despite the remarkable LC vascular robustness, IOP-induced distortions can potentially contribute to glaucomatous neuropathy.

## 1. Introduction

Vision loss in glaucoma is due to the loss of the retinal ganglion cells that transmit visual information to the brain.^1,2^ Glaucomatous axonal damage is believed to initiate within the lamina cribrosa (LC) region of the optic nerve head (ONH), where the axons exit the globe.^1–5^ Although neural tissue damage can occur even at normal levels of intraocular pressure (IOP), elevated IOP is a major risk factor for glaucoma and currently every method to prevent or treat glaucoma is based on reducing IOP.^6,7^ Nevertheless, the mechanisms by which elevated IOP contributes to the neuropathy remain unclear, complicating the development of new improved methods to prevent vision loss.^8–14^ A leading theory suggests that elevated IOP causes distortions of the ONH vasculature, particularly of the LC.^8,10,15–17^ The distorted vessels have reduced blood flow within, resulting in compromised perfusion and, most importantly, in reduced oxygenation of the neural tissues of the LC.^10,18,19^ Even mild reductions in oxygenation, if sustained, can result in or contribute to neural tissue damage.^20–23^

Despite remarkable advances of the experimental techniques available to study ocular biomechanics and blood flow, obtaining detailed maps of the LC flow and oxygenation remains out of reach. Available techniques suffer, for example, from limited penetration (OCT and related light-based techniques), ^24–28^ or insufficient spatial or temporal resolutions (ultrasound and MRI ^29,30^). Also importantly, several of these techniques are primarily aimed at detecting blood flow. While blood flow is important, studies note that it is crucial to measure and understand oxygenation.^27^ Oxygenation is related to, but not identical to flow. Techniques for measuring oxygenation within the blood vessels have been developed ^27^, but application in deep vessels within the LC remains out of reach. Because of the experimental challenges, several groups have turned to theoretical ^18,31^ and computational modeling ^9,10,32^ to study LC hemodynamics and oxygenation and get a better understanding of the role of IOP-induced deformations. Theoretical models are particularly elegant, but they are only able to account for highly simplified anatomies and thus are not yet useful to analyze the extremely complex nature of eye-specific vascular networks. Computational models have been able to overcome this limitation, first using 2D models, and later 3D flat LC anatomies.^9,10,18,33,34^

We have developed a technique to reconstruct eye-specific models of the complex 3D vascular network of the LC region ^35^ and then use it for hemodynamics and oxygenation simulation.^32^ Recently, we leveraged these techniques to study a set of eye-specific vascular networks and evaluate the effects of IOP-related tissue distortion on LC hemodynamics and oxygenation. We found that LC blood flow was sensitive to IOP-related vessel collapse, particularly if the collapses were clustered. An important finding was that normal local flow did not imply adequate oxygenation. In the study, however, we made several strong assumptions that we now aim to revisit in this work. For instance, as a first approximation we considered the vessels as either open and unaffected by IOP, or fully collapsed and closed. A better understanding of how IOP influences the LC, including blood flow and oxygenation requires a more precise and subtle consideration of how IOP distorts vessels, the flow within, and from these the tissue oxygenation.

Our goal in this study was to improve understanding of how LC oxygenation is affected by tissue deformations resulting from elevated IOP. We consider four eye-specific vascular networks from monkey ONHs. We analyzed the hemodynamics and oxygenation at baseline, as reconstructed, and then when subjected to experimentally-derived IOP-induced tissue deformations. Compared with our previous report, herein we use a more refined approach to map experimental tissue deformations into vessel deformations, which then alter LC oxygenation. Also, because of the simplistic assumption of how blood vessels were affected by IOP (open or closed) in our previous study, we were unable to evaluate details of the effects of IOP and we only considered baseline and high IOPs (10mmHg and 40 mmHg). Herein we used a much more realistic approach to estimate vessel deformations with IOP, and were thus able to obtain also a more refined understanding of tissue distortions and their effects.

## 2. Methods

*General procedure*. We reconstructed 3D models of the LC vessel networks of four healthy monkey eyes from histological sections following previously reported techniques.^32,35^ In-vivo IOP-related tissue deformations within the LC were determined from a healthy monkey using OCT imaging and analysis techniques under controlled IOP conditions ^36^ Two biomechanics-based techniques were then used to map the OCT-derived tissue strains to local LC to estimate their deformations. LC hemodynamics and oxygenations of the four vessel networks were simulated as reconstructed, baseline, and under several levels of IOP elevation (Fig 1). The outputs of these simulations were then analyzed to determine the effects of IOP on LC oxygenation (oxygen partial pressure, PO2). To interpret the results, the LC oxygen supply was classified into three levels based on the literature: normoxia (PO2 > 38 mmHg), mild hypoxia (38 mmHg > PO2 > 8 mmHg), and severe hypoxia (8 mmHg > PO2). Below we provide more details of each of the steps.

**Figure 1.**
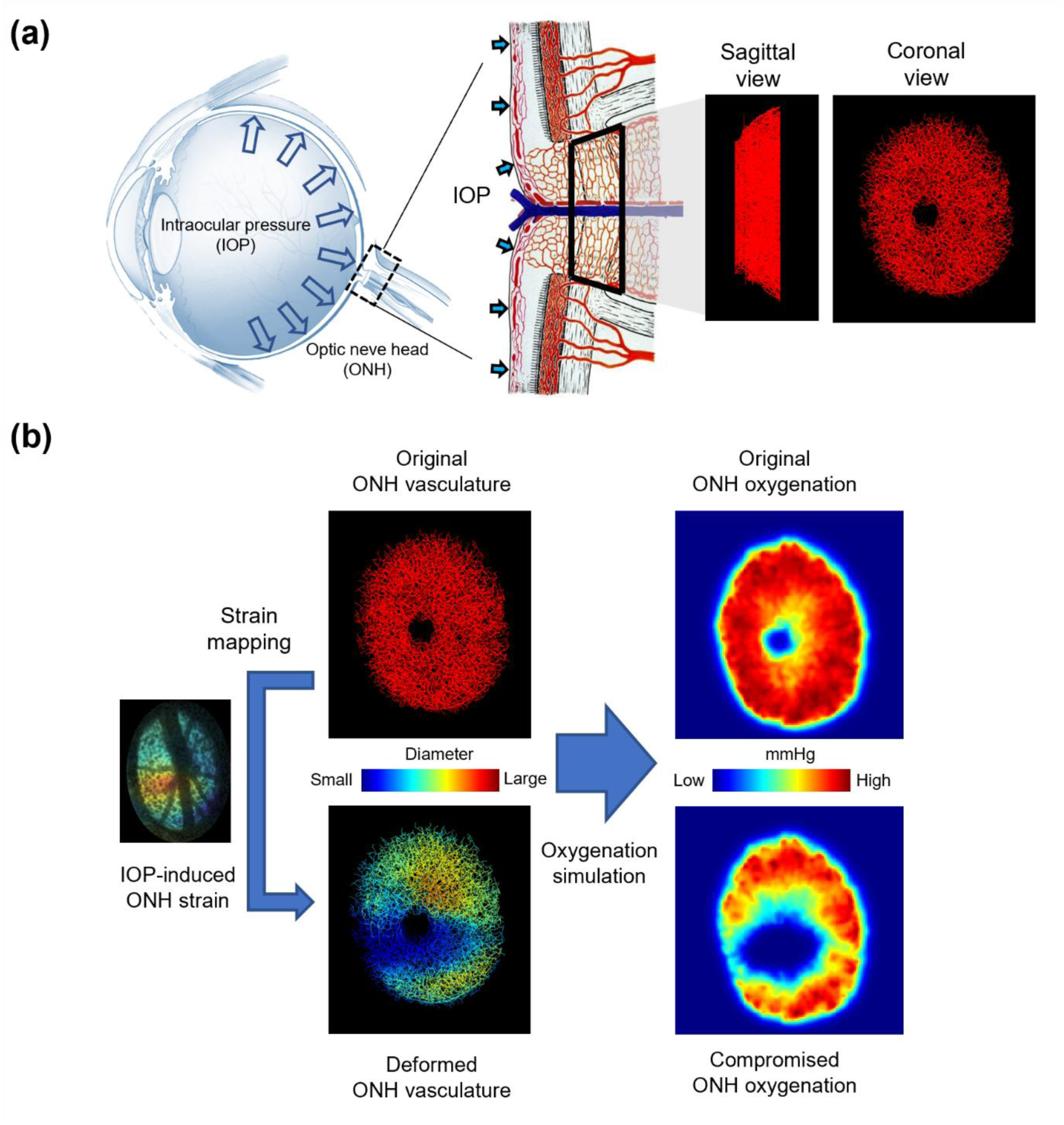
(a) Diagrams of the eye and ONH vasculature (left and middle). An eye-specific model of ONH vasculature (right). The region reconstructed is indicated by the black trapezoid in the center panel. The region is part of the scleral canal, delimited at the periphery by the connective tissues of the sclera and/or pia mater, at the center by the central retinal artery and vein and by flat planes perpendicular to the central retinal artery and vein, at locations in the pre-lamina and retro-lamina regions selected to ensure that the LC was completely enclosed. (b) Schematic of our analysis process. The four eye-specific ONH vasculatures were reconstructed from histological sections and their baseline hemodynamics and oxygenation predicted by simulation. Experimental IOP-induced 3D deformation maps were then applied and used to predict vessel segment-specific distortions. The full LC hemodynamics and oxygenation were then re-calculated for the networks with distorted vessels. The left hand-side of panel a was adapted from a diagram by NEI. The middle panel was adapted from a classic drawing. ^39^

### Vascular network reconstruction

All procedures were approved by the University of Pittsburgh’s Institutional Animal Care and Use Committee (IACUC) and adhered to both the guidelines set forth in the National Institute of Health’s Guide for the Care and Use of Laboratory Animals and the Association for Research in Vision and Ophthalmology (ARVO) statement for the use of animals in ophthalmic and vision research. The reconstructions were made following the processes described elsewhere.^32,35^ Briefly, four healthy female rhesus macaque monkeys’ heads and necks were processed for vessel labeling. The anterior chamber of each eye was cannulated to control IOP throughout the experiment. Polyimide micro-catheters were inserted into carotid arteries for vessel labeling, and warm phosphate-buffered saline (PBS) washed the vascular bed to remove blood. A lipophilic carbocyanine dye, Dil, was used to label the vessels in the eye. We perfused 100 ml Dil solution into each carotid artery, followed by a PBS wash, and then perfused 100 ml formalin into each carotid artery to fix the eye. Afterward, the eyes were enucleated, extraocular tissues were carefully removed and immersion fixed in 10% formalin for 24hrs to complete fixation. We used formalin fixation because it has been shown to only have a minimal effect on ocular tissues. ^37^

The ONH and surrounding sclera were isolated using a 14-mm diameter trephine, the tissues were then cryoprotected and cryosectioned (16 µm-thick) as described elsewhere.^32^ Fluorescence microscopy (FM) and polarized light microscopy (PLM) images were acquired to visualize vessels and collagen, respectively. Stacks of sequential FM and PLM images were aligned and registered in Avizo software (version 9.1). After registration, the FM images were segmented and skeletonized to create a 3D reconstruction of the ONH vasculature.^35^ Our reconstructed vascular networks included the whole LC region, and some of the pre-laminar and retro-laminar regions (Figure 2). This ensured that the 3D LC network was fully enclosed within the region reconstructed. Vessels in the LC region were identified based on the presence of collagen beams in PLM images. As in our previous work ^32^, we could not ensure that the vessel diameters in the ex-vivo sections were truly representative of the in-vivo condition, and therefore a uniform capillary diameter of 8 µm was assumed, as measured in ^38^.

**Figure 2.**
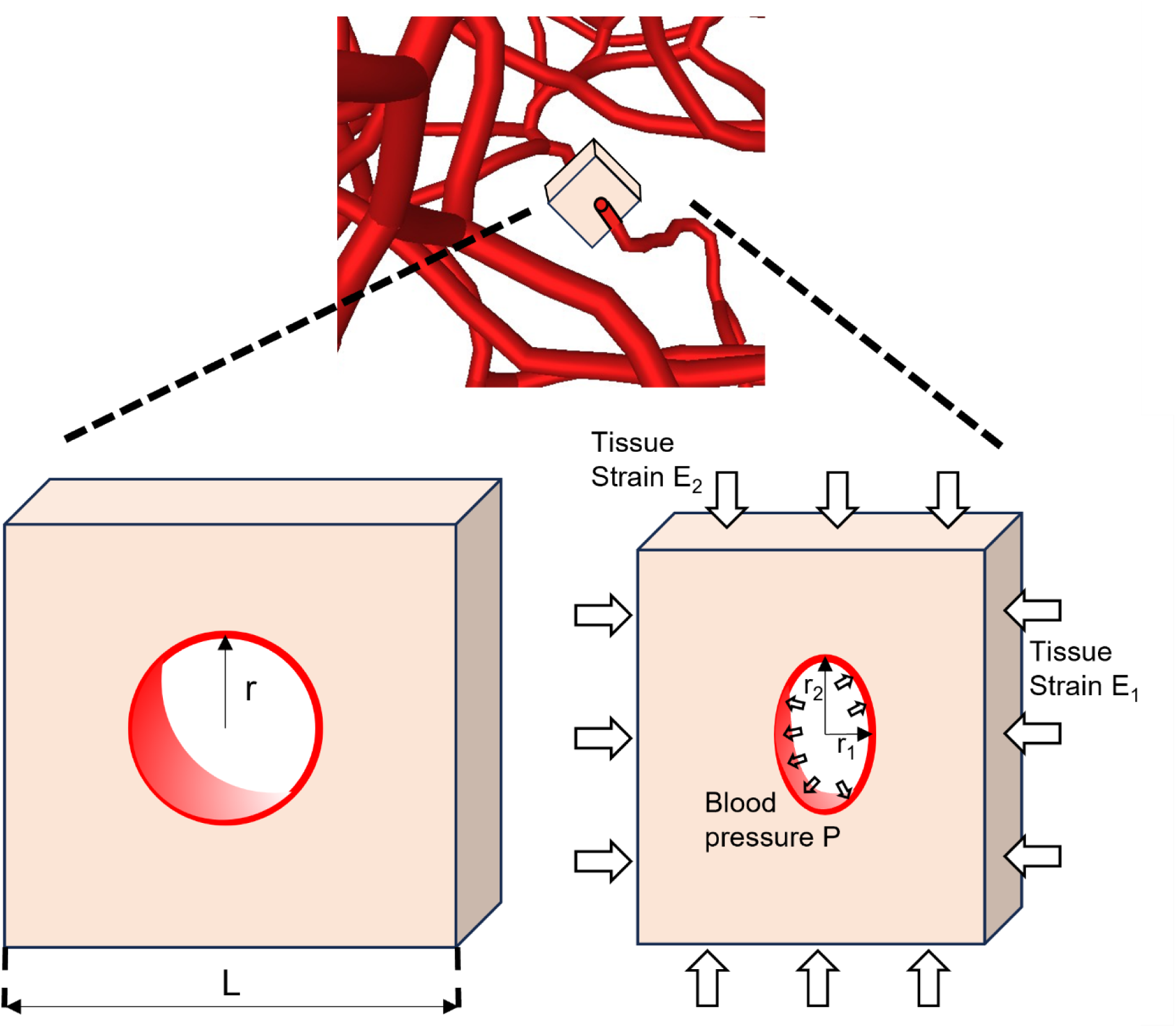
Diagram of the cross-sectional vessel deformation model. The vessel segment (red) was embedded in the center of the tissue (flesh color) with a square cross-section. The edge length L of the square was set to be 40 *μm*. The original radius (r) of the vessel was set to be 4 *μm*. The deformed vessel cross-section was considered as an ellipse with the semi axes of *r*_1_, *r*_2_.

### Vascular reconstruction validation

To minimize clotting, which could prevent dye perfusion from reaching all vessel segments, we did our best effort to minimize the time between animal death and perfusion, and extensively washed the ONH with PBS. We estimate that all perfusions were started less than 90 min from death. Using a large dye volume ensured sufficient labeling. FM revealed strong signals in retinal and choroidal vessels, indicating successful perfusion. In some regions signal was less strong, potentially due to partial occlusion of a vessel. However, these were still discernible by controlling the brightness and contrast settings.

Nevertheless, we acknowledge that there will always be some uneven labeling, discontinuities or leaks that could impact the visualization and reconstruction. Manual intervention including "cleaning" and "bridging" segments, was required. This could introduce potential artifacts and randomness in our final reconstructed model. To validate the reconstruction technique we reconstructed the same eye twice by two individuals working independently from the same sections. We then processed them in exactly the same way all the way to the computational predictions of hemodynamics and oxygen distribution.

### Vessel deformation

The deformation of blood vessels is a classic biomechanical process, which is determined by many factors, including the surrounding tissue strain and stress environment, the compliance of the vessel wall, and luminal blood pressure. ^40,41^ In this work, we assumed that the small vessels of the ONH were primarily deformed by the in-vivo ONH deformations (strain) resulting from an increase in IOP. This is the same general approach used elsewhere by us ^42^ and others.^10^ This work, however, differs from previous ones in that we used a more refined biomechanics-based method to translate the in-vivo ONH strains into local LC vessel deformations. Specifically, we performed in-vivo OCT scans on a health monkey while IOP was controlled at 10 mmHg(baseline), 20 mmHg, 30 mmHg, 40 mmHg, 50 mmHg, 60 mmHg. And a digital volume correlation (DVC) method was applied to measure the in-vivo ONH strain from the OCT volumes, as reported before. ^36^ Since the in-vivo ONH deformation and vascular networks are from different eyes, corresponding scaling and rotation were utilized to align the central retinal artery/vein and the temporal-nasal and superior-inferior axes. The approach was essentially the same as in a previous publication.^42^ We then used a biomechanics-based techniques to map the tissue strains to LC vessel distortions. These steps are described in more detail below.

### Deformation mapping

To map the local strain tensor to the local vessel segment we used an approach we call “cross-sectional strain mapping”. This approach is based on modeling the blood vessels as hollow tubes embedded in the ONH, with the blood pressure applied on the vessel wall. This approach has the advantage that the deformation of ONH vasculature takes into consideration the complex network geometry, hemodynamics, and vessel-tissue interaction. A fully coupled model, however, is extremely complex and difficult to simulate. We therefore took advantage of some simplifications to approximate vessel deformation and evaluate the changes in blood flow and oxygenation. The flow resistance of vessels, crucial in determining vascular network hemodynamics, is highly sensitive to their diameters (∝ *d*^4^). As a result, we focused on the cross-sectional deformation of each vessel, as it closely relates to diameter variations.

Each vessel segment was assumed to be embedded in a tissue with a square cross-section, with the vessel centerline is located at the center of the square (figure 2). The entire region consisted of tissue and the hollow vessel lumen. Here, we assumed that the vessel wall had the same material as the surrounding tissue. The boundary conditions of this system consist of: 1. The displacement boundary conditions on the side surface of the tissue region were interpolated from the in-vivo ONH tissue deformation tensor; 2. The blood pressure boundary condition applied on the interior vessel wall were derived from the hemodynamic simulation. We applied a linear elastic model to the behavior of the tissue, setting the stiffness (Young’s modulus) to 0.3 MPa and the Poisson’s ratio to 0.5. ^40^ Specifically, the deformation of the cross-section of the vessel satisfied the linear relationship:

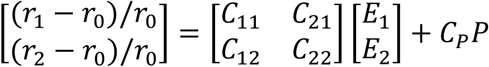

Where *r*_*x*_, *r*_*y*_ were the semi axes of the vessel cross-section, which was considered to be an ellipse under deformation, *r*_0_was the original radius of the vessel, *E*_*x*_, *E*_*y*_ were tissue strains added on the x, y direction, respectively, *P* was the luminal blood pressure (mmHg), and the coefficients *C*_11_, *C*_12_, *C*_21_, *C*_22_, *C*_*P*_ were 3.03, 0.38, 0.38, 3.03, 5.9 · 10^−3^, respectively. We generated models with various parameters in Abaqus software and simulated vessel deformations. Multilinear fitting techniques were then applied to derive the coefficients from the simulation results.

### Hemodynamic and oxygenation simulation

The hemodynamics of the four vascular networks were simulated at the baseline condition and subjected to the IOP-induced deformations as described in the previous section. The vascular network was represented as a system of interconnected capillary elements. Considering the low Reynolds number in capillaries, the blood flow inside a cylinder-like capillary can be approximately by the Poiseuille flow,

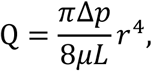

where Q is the blood flow rate, r is the vessel radius, L is the vessel length, µ is the blood viscosity, and Δ*p* is the pressure drop along the vessel. The blood viscosity µ was described as a function of vessel radius and hematocrit (i.e., the volume fraction of red blood cells). ^43,44^ It is worth noting that, both experimental observations and theoretical models reveal a dramatic increase in µ (viscosity) as the vessel diameter reaches the width of a red blood cell (∼2 µm).^44^ Namely, the vessel is nearly fully collapsed at that diameter. Flow and circulation of red blood cells in tight vessels can be extremely complex. Such analysis is beyond the scope of this work, but interested readers are encouraged to read the papers by Pries and Secomb mentioned above and by Ebrahimi and Bagchi. ^45^

When the vessel undergoes deformation due to IOP elevation, the cross-section of the vessel can be approximated as an ellipse, while the flow satisfies ^46^:

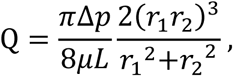

where the *r*_1_, *r*_2_ represent the axes of the ellipse cross-section of the vessel.

Pressure boundary conditions were applied to simulate the physiological blood supply for ONH. These were described and discussed in detail elsewhere.^42^ Specifically, we divided the model boundaries into four surfaces for assigning the blood pressure conditions that irrigate the ONH region. The arteriole inlet pressure, representing blood inflow from the circle of Zinn-Haller, was set to 50 mmHg at the periphery. The central retinal vein, responsible for drainage, had a venule outlet pressure of 15 mmHg. The anterior ONH boundary and the posterior ONH boundary were set to pressures of 20 mmHg and 16 mmHg, respectively.

The ONH oxygenation simulation relied on the blood flow field, which involved modeling the transport of oxygen from vascular networks to tissues, thereby establishing the oxygen field. The physical principles of oxygen transport are well established ^47^, including diffusion in the tissue and convection in the vessel. The governing equations for oxygen diffusion and consumption in tissue are:

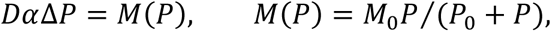

where *Dα* is the oxygen diffusion coefficient, and the oxygen solubility coefficient, respectively. *P* is the oxygen partial pressure in tissues. The oxygen consumption rate *M*(*P*) can be estimated by Michaelis-Menten enzyme kinetics, where *M*_0_, *P*_0_ are the Michaelis-Menten constants. In this study, *M*_0_, which represents the demand of neural tissues, was assumed to be uniform throughout the ONH.

The oxygen flux in blood vessels satisfies:

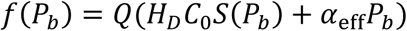

where *H*_*D*_ is the hematocrit, *C*_0_is the concentration of hemoglobin-bound oxygen in a fully saturated red blood cell, *P*_*b*_ is the blood oxygen concentration (mmHg), *S*(*P*_*b*_) is the oxygen-hemoglobin saturation as determined by Hill equation, and α_eff_ is the effective solubility of oxygen in plasma. All parameters are listed in Table 1.

**Table 1.** Parameters used in hemodynamic and oxygenation simulations.

| Hemodynamic parameters | Value | Reference |
| --- | --- | --- |
| Vessel diameter | 8 $\mu\text{m}$ | 32 |
| Arteriole pressure | 50 mmHg | 32 |
| Venule pressure | 15 mmHg | 32 |
| Anterior blood pressure | 20 mmHg | 32 |
| Posterior blood pressure | 16 mmHg | 32 |
| **Oxygenation parameters** |  |  |
| Oxygen diffusion coefficient, $D\alpha$ | $6 \times 10^{-10} \text{ mlO}_2/\text{cm/s/mmHg}$ | 55 |
| Effective oxygen solubility, $\alpha_{eff}$ | $3.1 \times 10^{-5} \text{ mlO}_2/\text{ml/mmHg}$ | 55 |
| Consumption rate, $M_0$ | $5 \times 10^{-4} \text{ mlO}_2/\text{ml/s}$ | 32 |
| Oxygenation at half-maximal consumption, $P_0$ | 10.5 mmHg | 55 |
| Maximal RBC oxygen concentration $C_0$ | 0.5 $\text{mlO}_2/\text{ml}$ | 55 |
| **Vessel deformation parameters** |  |  |
| Vessel compression parameter $C_{11}, C_{22}$ | 3.03 | See text |
| Vessel compression parameter $C_{12}, C_{21}$ | 0.38 | See text |
| Vessel compression parameter $C_p$ | $5.9 \times 10^{-3} / \text{mmHg}$ | See text |

We used a fast and efficient method to simulate the convective and diffusive oxygen transport in the complex ONH vascular networks, as reported before. ^48^ The comparison of oxygenation levels was analyzed between the baseline and deformed vessel networks to assess the impact of elevated IOP on the LC oxygenation.

### Hypoxia definition

We were interested in quantifying the effects of IOP on LC oxygenation. We therefore selected as outcome measure the size of the region suffering hypoxia, or the “hypoxia region” as the way to quantify the LC oxygen supply, and as an indicator of potential damage from the IOP elevation. Tissue hypoxia, characterized by an insufficient supply of oxygen to tissues, often arises from structural and functional disturbance in microcirculation. ^23,49^. Pathological processes driven by hypoxia involve complex biochemical mechanisms, and vary significantly across different tissues ^21,23,50^ A normal oxygen supply ensures physiological cellular activity, and a decrease in oxygenation increases the risk of damage to the tissues. When oxygen partial pressure declines to some critical levels, irreversible damage, such as tissue necrosis, can occur. ^51^ To distinguish the hypoxia level and the related potential tissue damage, we categorize tissues into three levels based on local oxygen partial pressure:

1. Normoxia (>38 mmHg): Normal cellular activity and metabolism.
2. Mild hypoxia (8∼38 mmHg): Physiological responses to hypoxia occur. If sustained chronically, may contribute to neural tissue damage.
3. Severe hypoxia (<8 mmHg): Tissue necrosis and irreversible damage at the short time scale.

In our previous study on LC oxygenation we provided a detailed rationale for our choices of oxygen tension levels.^42^ Briefly, 8 mmHg (∼1% oxygen) is a widely accepted threshold for severe hypoxia. ^20,50–54^ The threshold for normoxia, 38mmHg (∼5% oxygen), was chosen based on the literature. ^51^ Mild hypoxia is then defined as the intermediate state between normoxia and severe hypoxia. It is important to recall that this is not intended to suggest that there are no potentially ill effects of mild hypoxia. It seems plausible that the effects take longer or are more pronounced in tissues that are already in distress, for example due to mechanical insult. As outcome measures, we assessed the volumetric fraction of these three types of regions. The mild and severe hypoxia regions are mainly used to evaluate the LC oxygenation challenge caused by elevated IOP.

### Alternative approaches to mapping the compression

As noted above, there are currently no experimental methods suitable to measure IOP-induced distortions of the LC capillaries. Thus, it is necessary to define a way to map ONH strains to vessel segment distortions. Since this mapping directly determines the magnitude of the vessel distortions, it likely also plays a major role in the effects of IOP on the blood flow and oxygenation. While we still posit that the cross-sectional strain mapping technique described above is reasonable, we are aware that the vessel distortion is complex and may depend on many other factors and mechanisms. Therefore, there could be other ways to assign vessel segment distortion from the tissue deformation. We were interested in exploring if other strain mapping techniques could lead to very different effects on the blood flow and oxygenation. To evaluate this, we decided to pick an alternative method that would represent a realistic worst-case limit. For this, we considered an approach in which each vessel segment experiences an isotropic compression from the surrounding tissues. In this case the compression of the vessel radius was derived from the largest tissue compression, specifically, from the minimum eigenvalue of the strain tensor, *E*_3_. We called this approach “Isotropic compression mapping”. In the isotropic case, the cross-sectional equation turns into

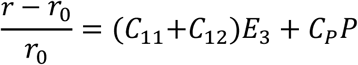

Where *r* and *r*_0_ are the compressed and original radius of the vessel, respectively. *E*_3_ is the maximum tissue compression. We repeated the whole set of simulations, including every strain mapping and simulation, using this alternative mapping approach. The rationale was based on the lack of direct experimental evidence that the cross-sectional mapping is accurate. In other words, we would anticipate that the actual effects of IOP would fall somewhere between the effects predicted using both approaches.

## 3. Results

### Reconstruction repeatability

The two vascular networks reconstructed independently from the same labeled eye exhibited extremely similar anatomy and oxygenation (Figure 3). Total vessel lengths were 547.68mm and 528.68 mm, a difference of 3.3%. The average LC oxygen partial pressures were 55.56 mmHg and 54.85 mmHg, a difference of 1.3%. Altogether this indicates excellent repeatability of our reconstruction process.

**Figure 3.**
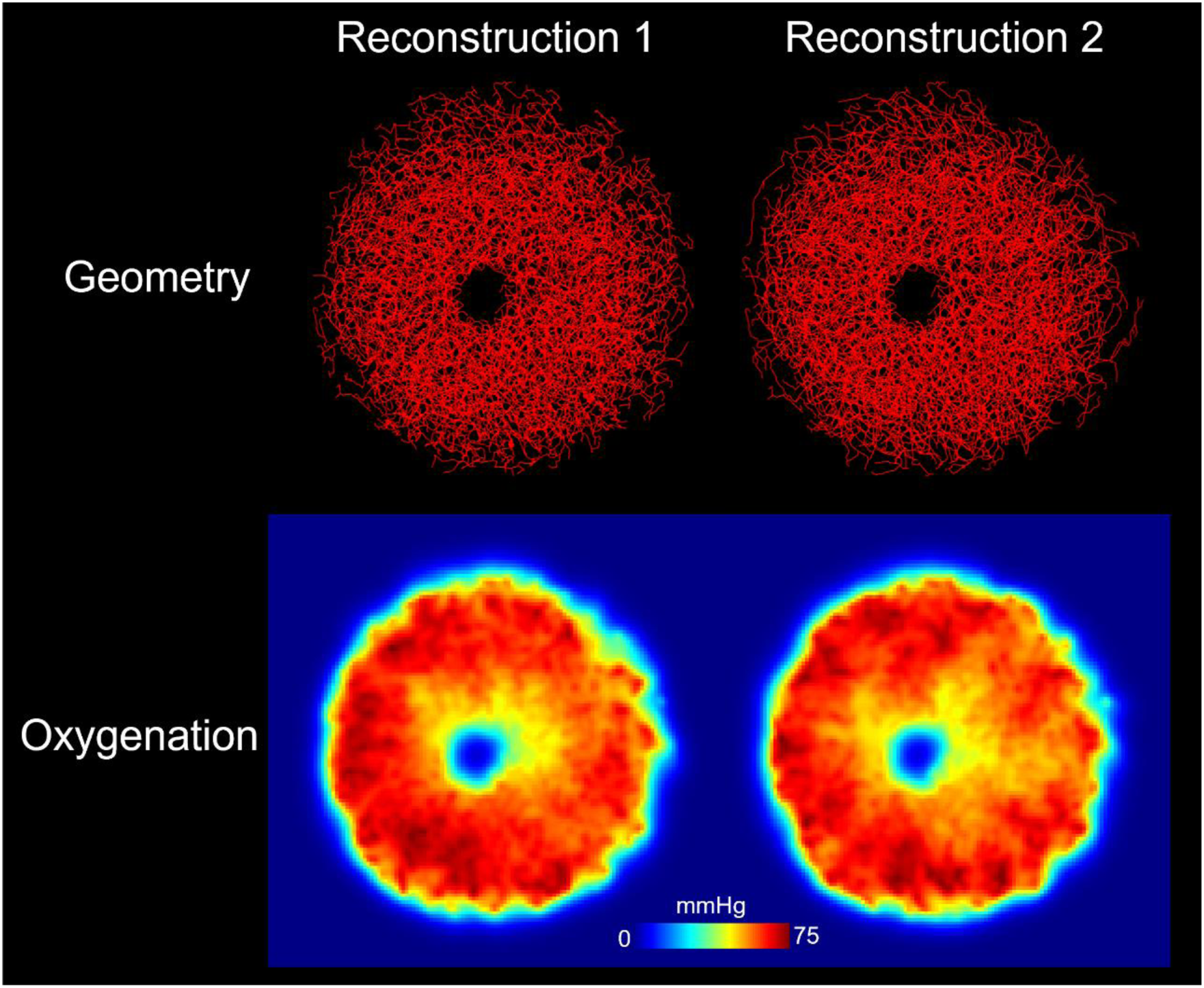
Two independent reconstructions from the same labeled eye. Both reconstructions exhibited similar geometry and oxygenation, indicting excellent repeatability in reconstruction.

### Four eye-specific reconstructions

The four vascular networks used in our simulations are shown in Figure 4. The DVC-based ONH deformations are shown in Figure 5.

**Figure 4.**
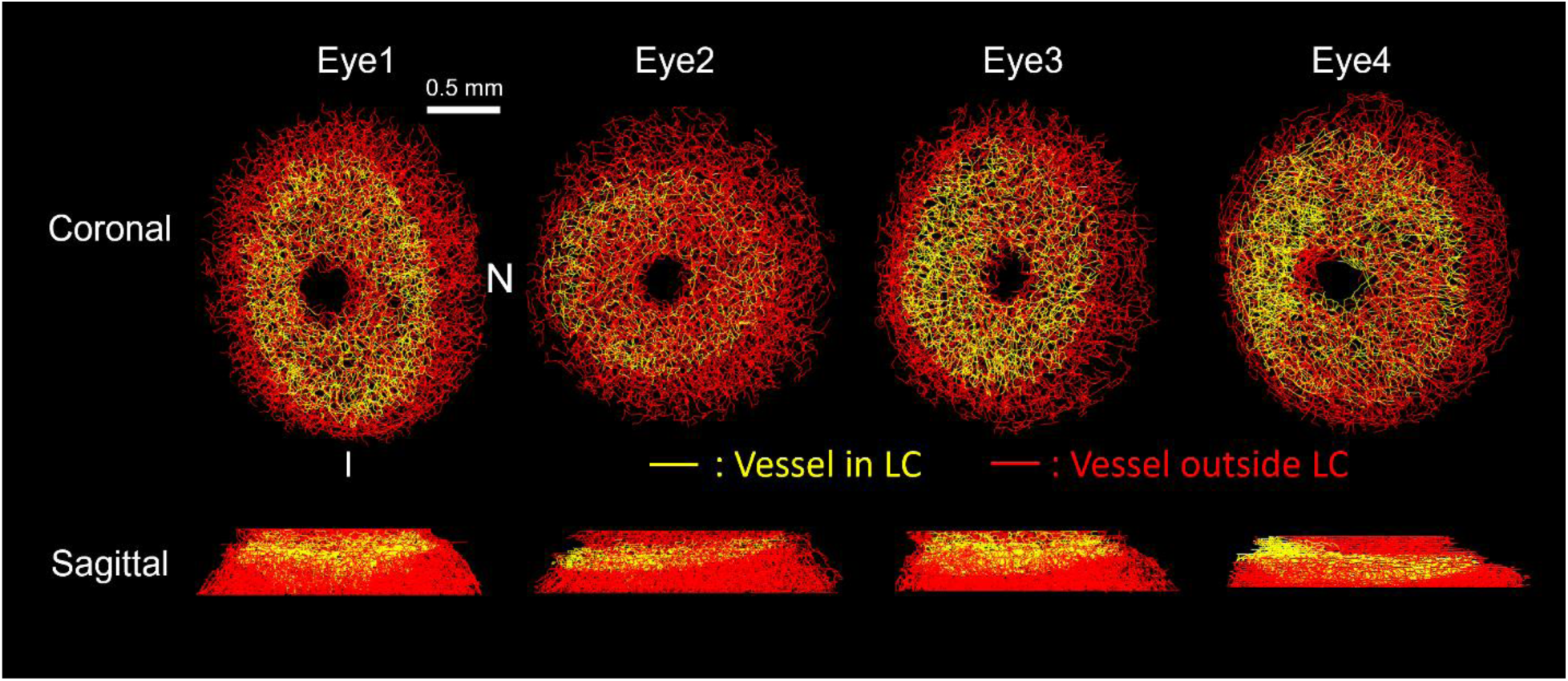
Reconstructed vasculatures of the ONH of the four eyes used in this work. All four networks were reconstructed from the right eyes (OD) of healthy monkeys. To improve boundary conditions we simulated the full volumes shown, but the analyses focused on the LC region (segments colored in yellow).

**Figure 5.**
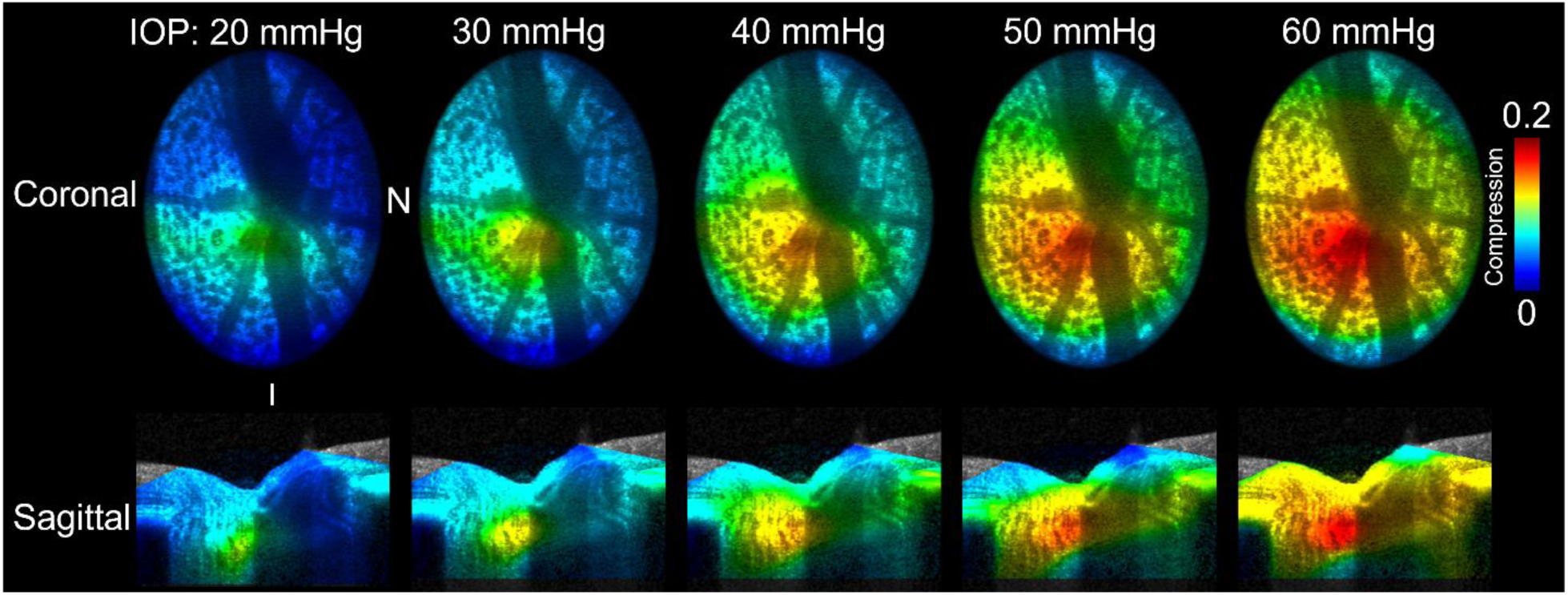
The ONH compression fields under various IOPs. The fields were derived using DVC technique to quantify tissue deformation from OCT images acquired at the baseline (10 mmHg) and elevated IOPs. The magnitude of the compression increased with IOP. The coronal views of deformations are shown in OD configuration. Note that colors indicating compression are only shown where the compression was measured with the DVC. A DVC region was defined that completely enclosed the scleral canal region where the vasculature was modeled to ensure accurate compressions were computed everywhere necessary.

### Effects of IOP-induced tissue distortions

The effects of IOP increases on LC oxygenation using the cross-sectional approach are shown in Figures 6-8. The detailed vasculature deformation and oxygenation of Eye 1 are shown in Figure 6. At baseline all vessel segments were assumed to be 8 µm in diameter. As IOP increased, tissue distortions led to decreased vasculature diameters, and with this to also decreased ONH oxygenation. For briefness, we only show one eye. Similar patterns were observed for the four other eyes.

**Figure 6.**
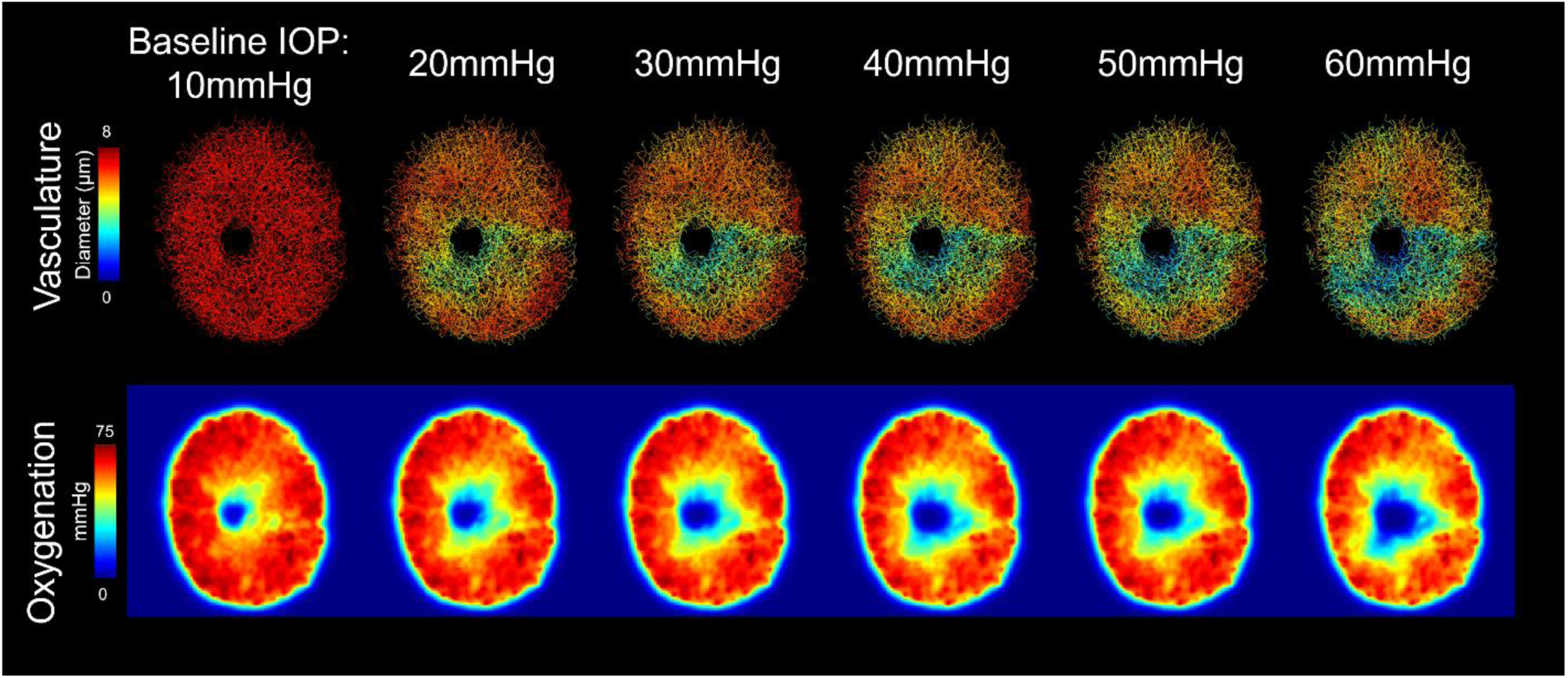
Eye1 vasculature deformation and oxygenation at different IOPs for cross-sectional strain mapping. The diameters of the vasculature and the oxygenation decreased with IOP elevation. Regions of low oxygenation are primarily near the central portion of the ONH, although it appears to be more evenly split between the superior and inferior portions of the disc than one might assume given that the distortion and vessel compression seems concentrated in the inferior region. High IOP (>40 mmHg) caused a substantial region of vessel deformation (green and blue). As we mentioned in method section, vessel would nearly fully collapse when its diameter reaches the RBC’s size (∼2 µm). Here, the vessel diameters never reach that value.

The LC oxygenation frequency distribution and the hypoxia region under various IOPs are shown in Figure 7, 8. The IOP-induced deformations reduced LC oxygenation significantly in all the eyes (p<1e-5). The effects of IOP were largest with the IOP change from 10 to 20 mmHg. Further IOP increase had smaller effects per unit of IOP increase.. The IOP elevation primarily contributed to the expansion of the mild hypoxia region, whereas the severe hypoxia region remained small (single digits %), even in cases of extremely high IOP. The effects of IOP varied between eyes (different panels in Figure 7 and different symbols in Figure 8), but the trends were similar.

**Figure 7.**
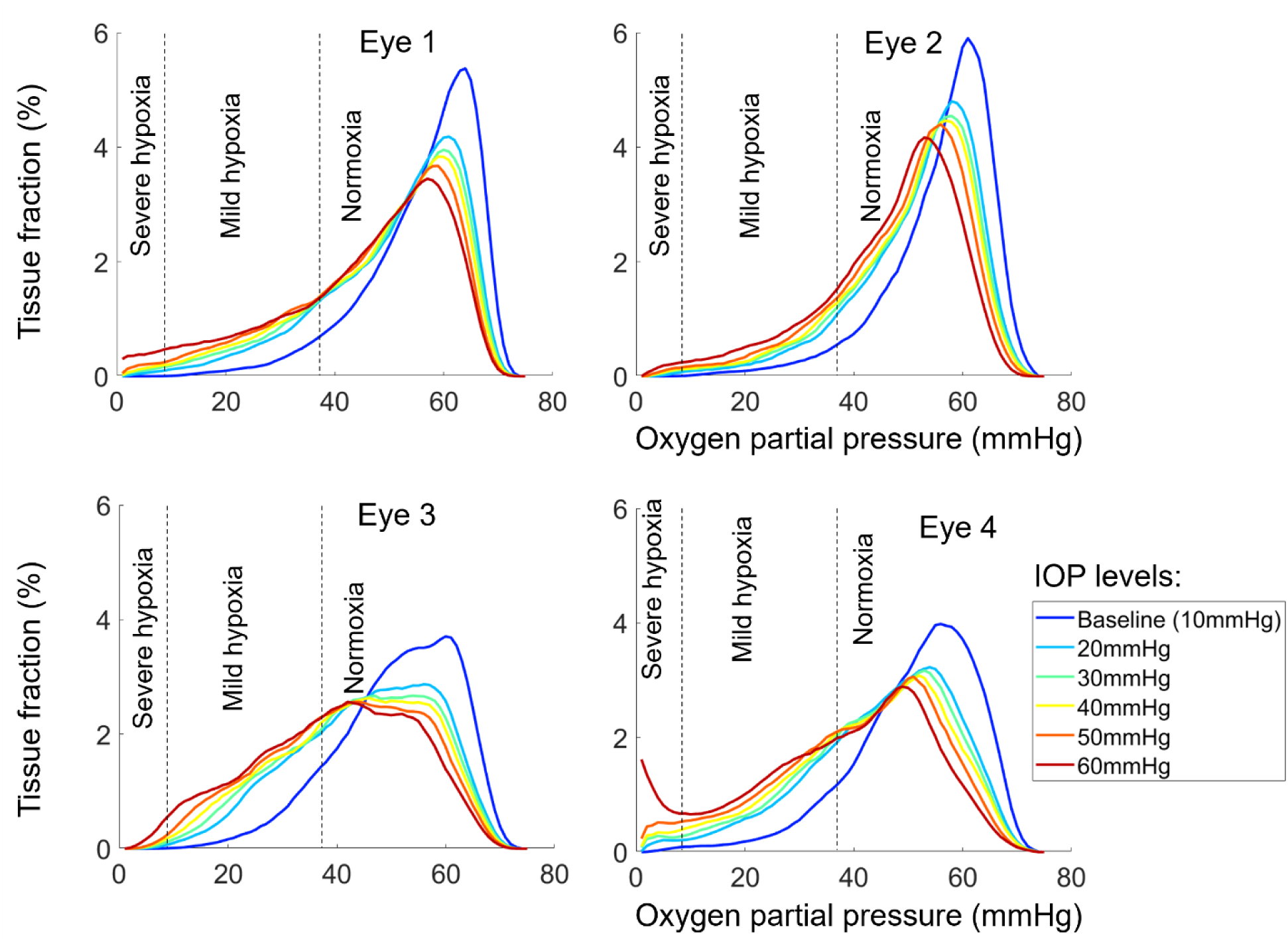
LC oxygenation frequency distribution for the four eyes at six IOPs under cross-sectional deformation mapping model. The oxygen frequency curves granually shift to the left as IOP increases for all four eyes. This indicates an association between elevated IOP and decreased oxygenation within the LC. The most substantial change in LC oxygenation frequency is observed during the initial rise in IOP (Baseline-20 mmHg). On the other hand, even under high IOP, only a minimal proportion of tissue falls under severe hypoxia.

**Figure 8.**
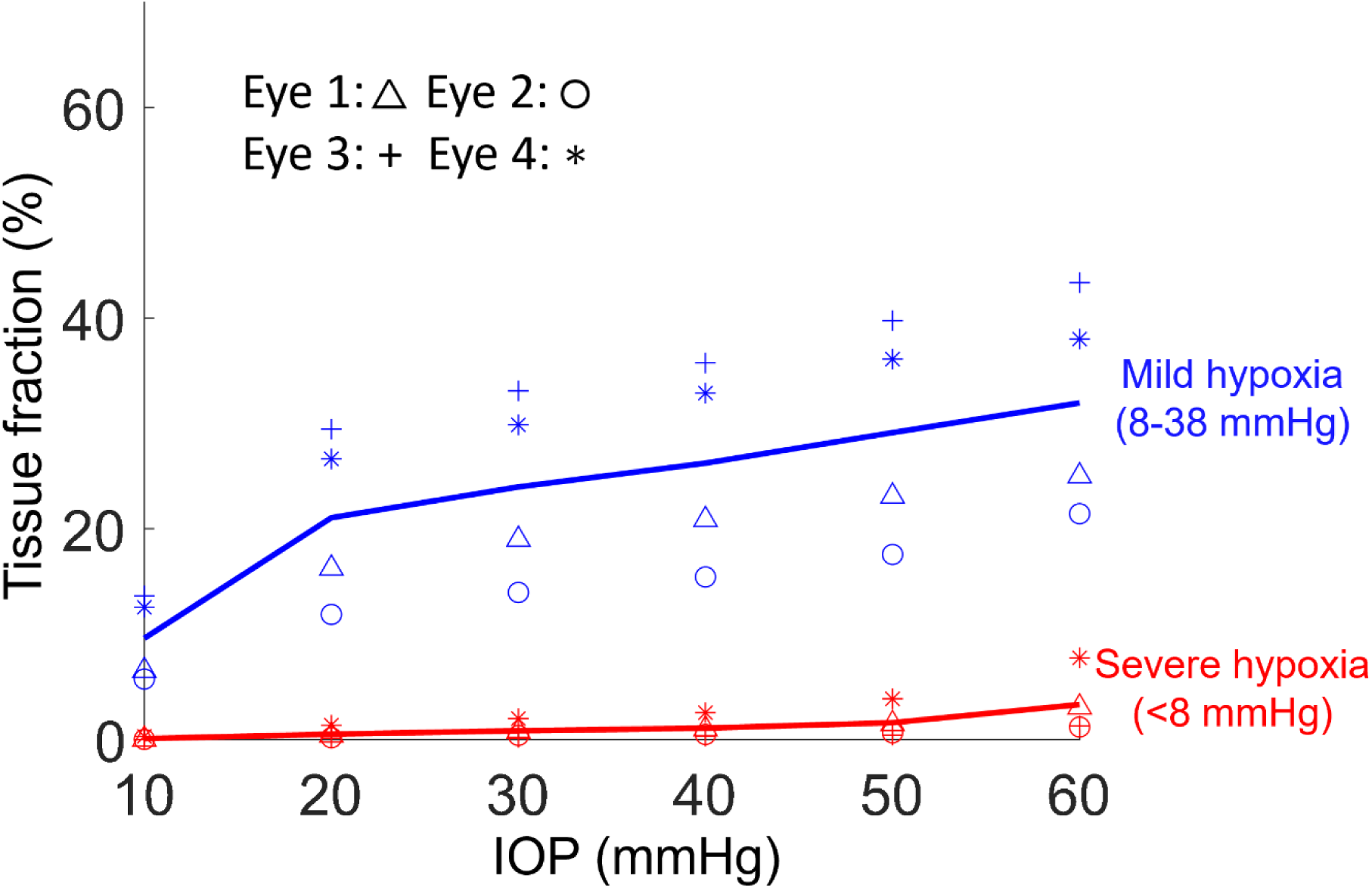
The relationship between IOP and hypoxia region in LC under a cross-sectional model. Results varied slightly between eyes, but overall, both the mild hypoxia and severe hypoxia region increased as IOP increased. For moderate IOP elevation (20∼30 mmHg), the mild hypoxia region increased significantly (p<0.05). About 25% of LC tissue suffered from mild hypoxia when IOP reached 30 mmHg. Only few severe hypoxia regions (<5%) were observed under extreme high IOP (>50 mmHg).

As explained in the Methods, we considered an alternative isotropic compression mapping method for mapping the DVC-measured deformations caused by IOP increases onto vessel distortions. The results were similar in distribution to those shown above using the cross-sectional mapping method, except that the effects were larger. The most substantial difference was that vessel compression did result in a substantial region under severe hypoxia (<8 mm Hg). The size of this region reached up to 40% at 60 mmHg in one eye. For briefness these results are presented as Supplementary material.

## 4. Discussion

Our goal was to evaluate how LC oxygenation is affected by the tissue distortions resulting from elevated IOP. Specifically, we used 3D eye-specific numerical models of the LC vasculature subjected to experimentally-determined IOP-induced tissue deformations for several IOP levels from 10 mmHg to 60 mmHg. DVC-determined tissue strain was mapped to vascular deformations using first a cross-sectional strain mapping technique and later an isotropic compression technique. We then used numerical simulations to determine LC hemodynamics in each case, and from these the oxygenation throughout the LC, with particular attention to measure the fraction of LC under mild or severe hypoxia. Three main findings arise from the models in this work. **First**, LC oxygenation was generally predicted to decrease as IOP increases. **Second,** moderately elevated IOP (20-30 mmHg) can lead to mild hypoxia in a substantial part (>20%) of the LC. Severe hypoxia region occurred for extreme IOP elevations (50-60 mmHg). **Third**, although IOP-induced deformations were roughly proportional to the level of IOP increase, the effects on LC oxygenation were larger at the initial increases in IOP, decreasing for higher IOPs. Below we discuss in detail each of these results, as well as the most important aspects of our modeling that readers should keep in mind when interpreting our results.

### LC oxygenation generally decreased as IOP increased

From baseline (10 mmHg) to extremely elevated IOP (60 mmHg), the oxygen distribution curve showed a general decreasing trend with IOP elevation. The hypoxic regions, both mild and severe, enlarged with IOP increases.

The association between LC oxygenation and IOP can be explained as follows. Qualitatively, the entire ONH is deformed during an IOP elevation. The fragile vessels within experience stretch, compression and distortion. A deformed vasculature will then alter the LC microcirculation, and potentially compromise the LC blood/oxygen supply. In this work, from quantitative perspective, we found that even a small IOP-induced ONH deformation (e.g. ∼5% compression under 30 mmHg of IOP) leads to a substantial decrease in LC oxygenation (∼25% mild hypoxia). Our previous work ^42^ also showed that LC oxygenation is susceptible to IOP-induced deformation. This can be attributed to two main reasons. First, the capillaries are softer and more susceptible to deformations than the fiber-rich tissues in the LC. (Camasão and Mantovani, 2021) The cross-sectional strain mapping model shows that the deformation of a capillary is approximately three times that of the surrounding tissue. Therefore, elevated IOP will induce a larger deformation on the LC vessels, leading to smaller diameters. Second, the flow conductance, defined by the ease with which blood can flow through, is sensitive to the vessel diameter, primarily from the quartic power relationship in Poiseuille’s law, and the diameter-dependent blood viscosity for red blood cells. (Pries and Secomb, 2008) These two effects amplify the influence of IOP on LC blood supply, and consequently, on LC oxygen supply.

### Moderately elevated IOP (20-30 mmHg) can lead to mild hypoxia in a substantial part (>20%) of the LC. Some severe hypoxia regions occurred for extreme IOP elevations (50-60 mmHg)

IOP-induced ONH deformation increased with higher IOP levels, resulting in a significant expansion of the LC hypoxia region. Based on the physiological and pathological scenarios, mild hypoxia (<5% oxygen) and severe hypoxia (<1% oxygen) were selected as indicators to evaluate the potential damage from the elevated IOP on LC oxygenation.

Mild hypoxia is considered to occur at oxygenation levels that hypoxia responses start being detected. ^51^ Short-term mild hypoxia can be reversed by the blood flow autoregulation.^56,57^ However, if mild hypoxia is sustained chronically, it may contribute to neural tissue damage. Therefore, our models predict that long-term, moderately elevated IOP may lead to progressive and irreversible neural tissue effects, potentially resulting or contributing to glaucoma.

Severe hypoxia can cause immediate and irreversible tissue damage. Individuals experiencing acutely elevated IOP are at risk of rapid vision loss and neural tissue damage ^58^, This could be seen as a contradiction: only a few severe hypoxia regions (<5%) were observed under extreme high IOP, even though experimentally such a high IOP level will likely lead to widespread damage in a short time.^59^ One potential explanation could be that the extreme high IOP could impact eye circulation and vision signaling other than the lamina region. For example, an acute extremely high IOP challenge can damage retinal circulation in closed-angle glaucoma.^58^ We focus on the effects of milder, chronically elevated IOP on the LC region, and therefore could underestimate the effects in extreme cases. Conversely, moderately elevated IOP typically does not lead to sudden vision loss or acute neural tissue damage. This is consistent with our findings that severe hypoxia does not occur in moderately elevated IOP.

It is worth noting that our results rely on the local vessel deformation mapping method. Because the actual vessel deformations resulting from IOP are still not known, we used two methods, cross-sectional strain mapping to and isotropic compression mapping as reasonable estimates. Although the trends were similar, the IOP effects were larger in the isotropic mapping case, with larger mild/severe hypoxia regions. Further experimental and modeling work is necessary to address the IOP effect on LC oxygenation and provide more clinical insight.

### Although IOP-induced deformations were roughly proportional to the level of IOP increase, the effects on LC oxygenation were larger at the initial increases in IOP, decreasing for higher IOPs

As IOP increased, ONH deformations became more pronounced. This deformation compromises the blood supply in ONH, and results in reduced LC oxygenation. Interestingly, the most significant change in LC oxygenation occurred as IOP began to rise from 10 mmHg to 20mmHg. This can be understood through the IOP-induced deformation and the compliance of the LC, defined as the ability of the LC to deform in response to pressure changes. Experimental observations have indicated that LC compliance exhibits a nonlinear behavior in response to elevated IOP. ^60^ When IOP experiences a moderate increase, meaning it deforms more easily. This higher compliance at moderate IOP levels results in rapid and substantial deformation of the LC.

As IOP continues to rise beyond moderate levels, the LC becomes less compliant, and its deformation grows at a slower rate. This indicates that the LC stiffens with higher pressure, making it less capable of further deformation. This nonlinear compliance behavior explains why the most substantial reduction in LC oxygenation occurs during the early stages of IOP elevation.

Additionally, this finding is consistent with the crimp behavior of LC fibers. Crimping refers to the wavy pattern of collagen fibers in the LC. When subjected to moderate IOP increases, these crimped fibers can stretch and straighten, contributing to the initial higher compliance and significant deformation. As the fibers straighten out, the LC becomes less compliant, leading to a slower growth of deformation with further increases in IOP (Foong et al., 2023).

This study integrates 3D eye-specific LC vascular networks with in-vivo ONH deformation data. We further employed a biomechanical-based strain mapping approach to capture vessel deformations. This allowed us to perform a more realistic hemodynamic and oxygenation simulation for elevated IOP compared to highly simplified generic cases, and thus providing a more precise evaluation of the IOP impact. Additionally, our work provides a systematic analysis of multiple eyes with low to high IOPs. The use of multiple eye-specific vascular networks reduced the influence of individual variances, enhancing the robustness of our findings.

It is important to consider the limitations of this study. We did not use vascular networks and ONH compression in our analysis. Both the vascular network and tissue deformation have significant individual differences. We applied scaling and rotation to align the central retina vessels and the temporal-nasal-superior-inferior axis for them.

Another limitation is that our work only considers the static status of LC hemodynamic and oxygenation. However, in-vivo blood/oxygen supply involved various regulation mechanisms to meet the changing demand of organisms. Short-term blood flow autoregulation has been identified as a significant factor in hemodynamics and oxygenation for eye. ^57,61^ Long-term vessel remodeling in the LC was also observed in the development of glaucoma.^5,62^ Although the precise regulation and remodeling in LC remain unknown, we acknowledge that they could alter the LC blood and oxygen supply. ^57,61,63–66^ Future research should incorporate the dynamic aspects of the LC blood and oxygen supply, which could contribute to the development of some pathological scenarios, such as glaucoma.

We employed cross-sectional strain mapping methods. While we considered the luminal vessels embedded within the tissue, our approach simplified the model by assuming the vessel wall and the fiber-rich LC tissue as homogeneous linear elastic materials. This simplification serves two primary purposes: Firstly, it helps to reduce the computational cost of the coupled system. Secondly, it avoids the need for certain parameters in a more complex non-linear model, which is hard to measure in experiments. Based on the homogeneous linear assumption, our method may under/over-estimate the vessel compressions in the large deformation case. Also importantly, our strain mapping approaches considered the detailed eye-specific vascular network, but not the corresponding eye-specific collagenous network of beams and pores. Considering the recent findings that the vascular and connective tissue networks of the LC are distinct^67^, could cause a much more complex mechanical interplay than what can be captured by the modeling in this work. Future studies should strive to evaluate the potential effects of incorporating both specimen-specific vessels and specimen-specific LC beams. Note that the above does not mean that we have ignored the LC connective tissues, it is just that their role is incorporated indirectly through the use of experimentally-derived deformation maps.

In summary, we analyzed the LC oxygenation under various elevated IOP cases. Moderately elevated IOP can lead to mild hypoxia in a substantial part of the LC, which, if sustained chronically, may contribute to neural tissue damage. For extreme IOP elevations, severe hypoxia was predicted, potentially causing more immediate damage. The findings provide a systematic picture of the IOP-induced deformation effect on the LC, including multiple LC vascular networks, two strain mapping techniques and ONH deformation under various IOPs. This is important to help understand ONH physiology and pathology.

## 6. Appendix

### Supplementary material

#### Effects of IOP increases on LC oxygenation obtained usign the isotropic compression mapping

The impact of IOP variations on oxygenation for the isotropic compression model are shown in Figures S1 to S3. The detailed vasculature deformation and oxygenation of Eye 1 are shown in Figure S1. To simplify comparison with the cross-sectional mapping approach the three figures in S1, S2 and S3 were prepared in the same way as figures 6 to 8. Similar to the cross-sectional case, as IOP increases, tissue distortions lead to decreased vasculature diameters, and with this to also decreased ONH oxygenation. However, the vessel deformation and the LC oxygenation of the isotropic compression case are overall larger than the cross-sectional case.

The LC oxygenation frequency distribution and the hypoxia region under various IOP for the isotropic compression model are shown in Figure S2 and S3. Compared to the cross-sectional case, the IOP-induced deformations caused to a larger LC oxygenation decrease. Moderately elevated IOP (20-30 mmHg) can lead to mild hypoxia in a substantial part (∼40%) of the LC. Severe hypoxia was predicted for extreme IOP elevations (>50 mmHg).

**Figure S1.**
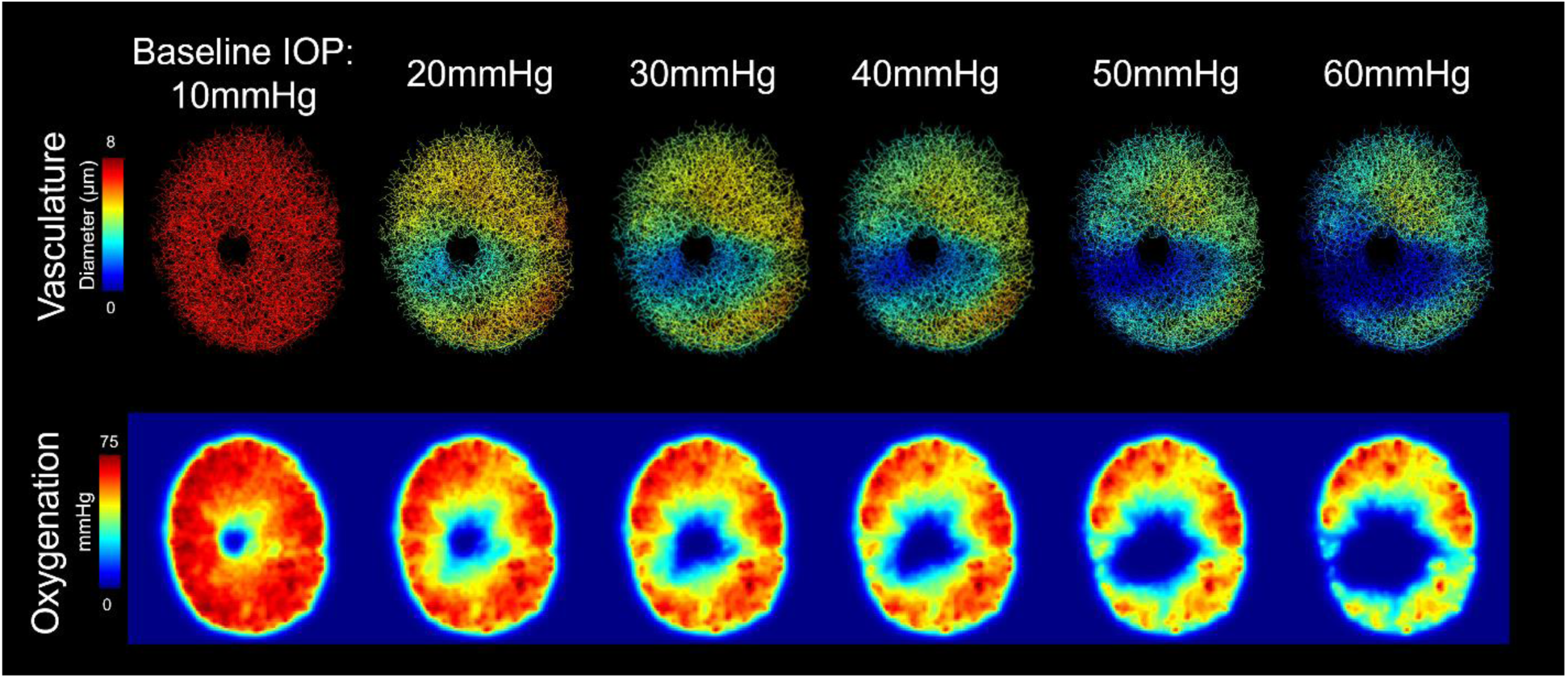
Eye1 vasculature deformation and oxygenation at different IOPs for isotropic compression strain mapping. The diameters of the vasculature and the oxygenation both decrease with IOP increases. Regions of low oxygenation and vessels with large compression had similar spatial distributions. High IOP (>40 mmHg) induced substantial vessel deformation (green and blue region), including the region of collapsed vessel (∼0 diameter).

**Figure S2.**
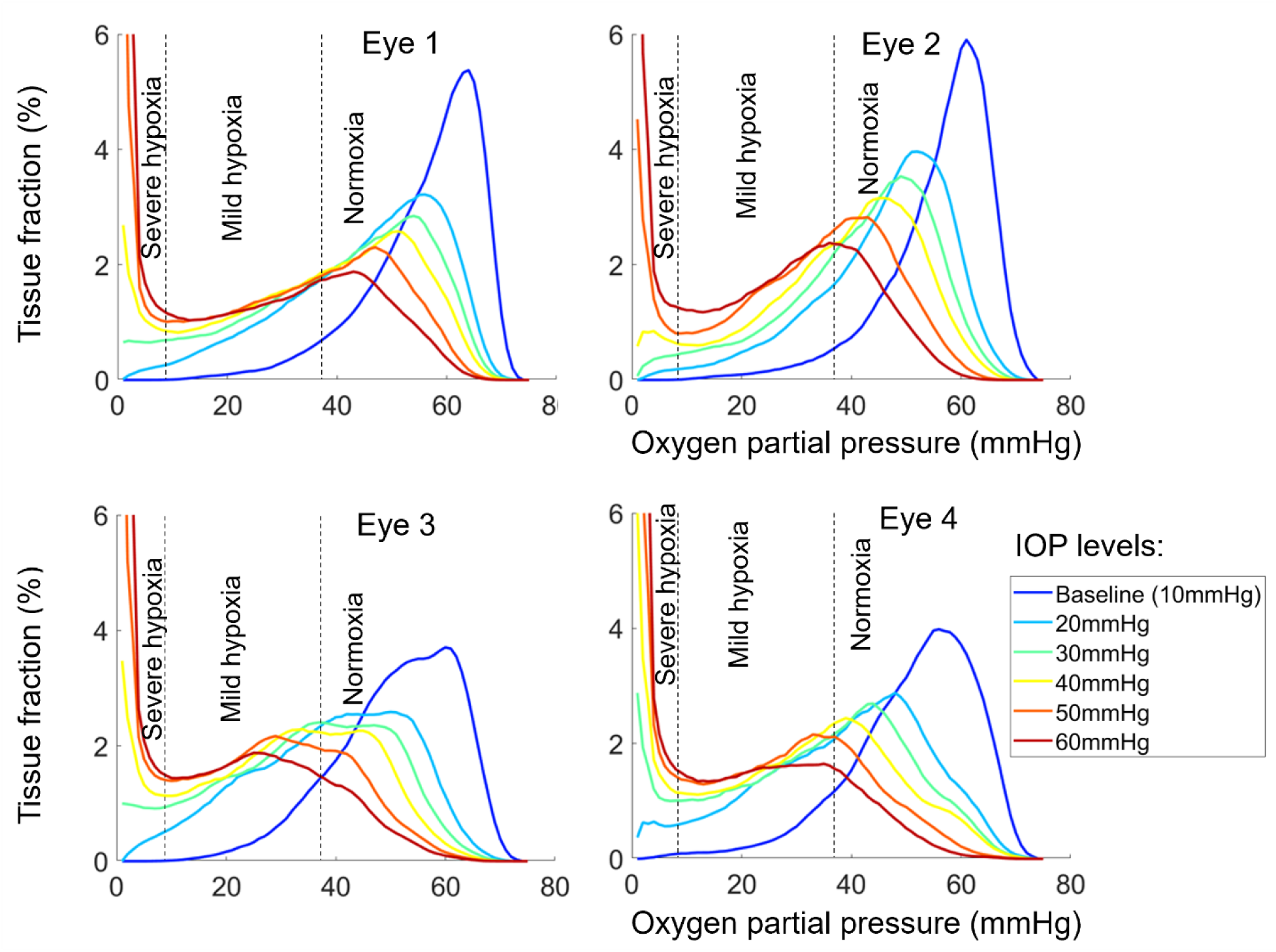
LC oxygenation frequency distribution for the four eyes at six IOPs under isotropic compression model. As IOP increases, the oxygen frequency curves granually shift to the left, indicating the decrease of the LC oxygenation. Meanwhile, the left side of the curves arise, indicating the increase in the area of severe hypoxia. Similar to the cross-sectional case, the most substantial change in LC oxygenation frequency is observed during the initial rise in IOP (Baseline-20 mmHg).

**Figure S3.**
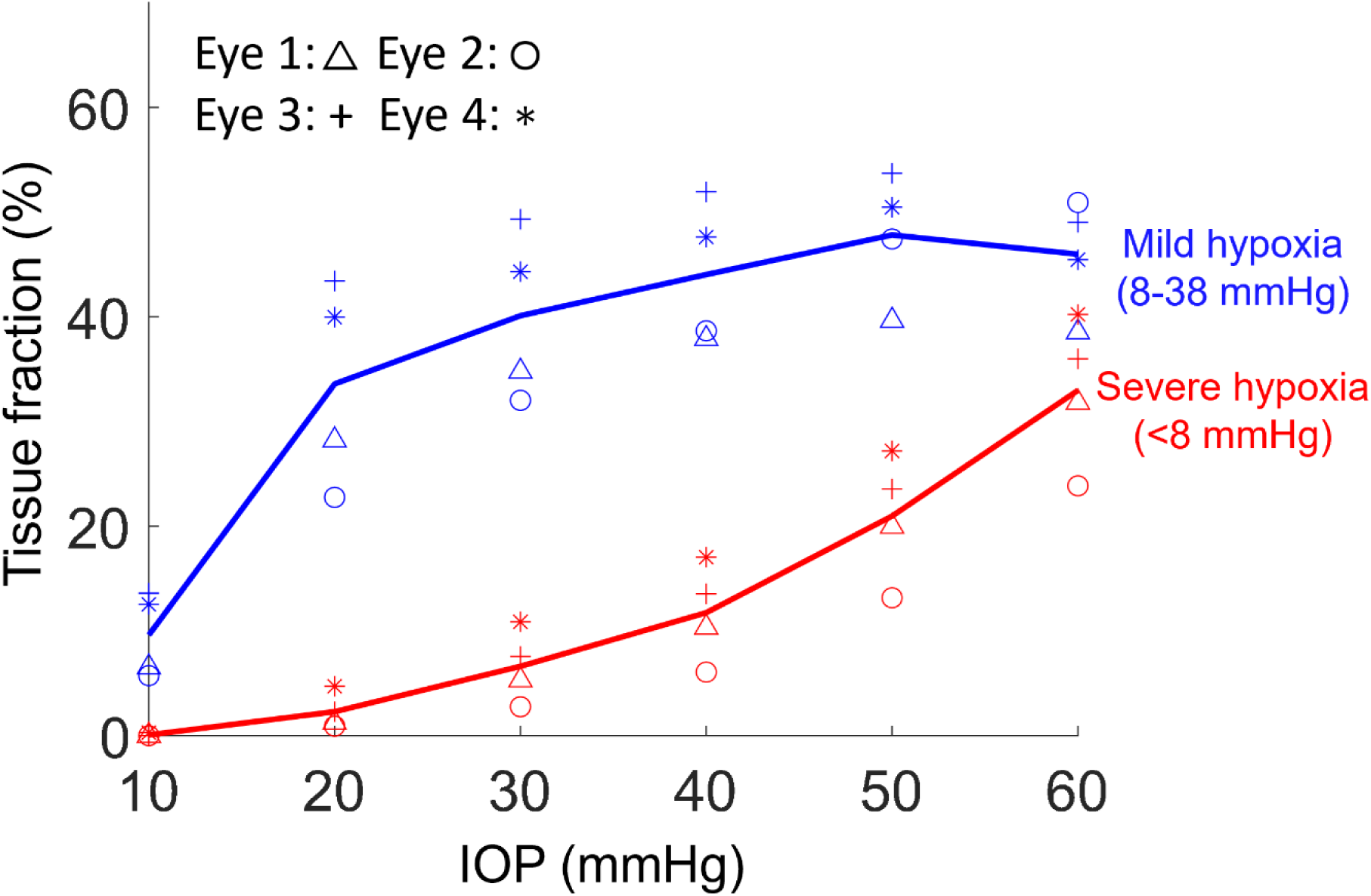
Statistic results for the relationship between IOP and hypoxia region in LC under isotropic compression model. For all four eyes, LC tissue fractions at mild and severe hypoxia increased with IOP. For moderate IOP elevation (20∼30 mmHg), the region of severe hypoxia was small, but the mild hypoxia region increased significantly (p<0.05). About 40% of LC tissue suffered from mild hypoxia when IOP reached 30 mmHg. Extreme IOP increases (>50mmHg) led to substantial severe hypoxia regions (>20% LC).

## References

1. Quigley H, Anderson DR. The dynamics and location of axonal transport blockade by acute intraocular pressure elevation in primate optic nerve. Investigative ophthalmology & visual science. 1976;15(8):606–616.

2. Nickells RW. The Cell and Molecular Biology of Glaucoma: Mechanisms of Retinal Ganglion Cell Death. Investigative Ophthalmology & Visual Science. 2012;53(5):2476–2481. doi:10.1167/iovs.12-9483h

3. Howell GR, Libby RT, Jakobs TC, et al. Axons of retinal ganglion cells are insulted in the optic nerve early in DBA/2J glaucoma. The Journal of cell biology. 2007;179(7):1523–1537.

4. Quigley H. vol. 377. Glaucoma Lancet. 2011:1367–1377.

5. Quigley HA, Nickells RW, Kerrigan LA, Pease ME, Thibault DJ, Zack DJ. Retinal ganglion cell death in experimental glaucoma and after axotomy occurs by apoptosis. Investigative ophthalmology & visual science. 1995;36(5):774–786.

6. Pitha I, Du L, Nguyen T, Quigley H. IOP and glaucoma damage: The essential role of optic nerve head and retinal mechanosensors. Progress in Retinal and Eye Research. 2023:101232.

7. Stowell C, Burgoyne CF, Tamm ER, et al. Biomechanical aspects of axonal damage in glaucoma: A brief review. Experimental eye research. 2017;157:13–19.

8. Burgoyne CF, Downs JC, Bellezza AJ, Suh J-KF, Hart RT. The optic nerve head as a biomechanical structure: a new paradigm for understanding the role of IOP-related stress and strain in the pathophysiology of glaucomatous optic nerve head damage. Progress in retinal and eye research. 2005;24(1):39–73.

9. Chuangsuwanich T, Birgersson KE, Thiery A, Thakku SG, Leo HL, Girard MJ. Factors influencing lamina cribrosa microcapillary hemodynamics and oxygen concentrations. Investigative ophthalmology & visual science. 2016;57(14):6167–6179.

10. Chuangsuwanich T, Hung PT, Wang X, et al. Morphometric, hemodynamic, and biomechanical factors influencing blood flow and oxygen concentration in the human lamina cribrosa. Investigative ophthalmology & visual science. 2020;61(4):3–3.

11. Sigal IA. Interactions between geometry and mechanical properties on the optic nerve head. Investigative ophthalmology & visual science. 2009;50(6):2785–2795.

12. Sigal IA, Grimm JL. A few good responses: which mechanical effects of IOP on the ONH to study? Investigative ophthalmology & visual science. 2012;53(7):4270–4278.

13. Voorhees AP, Hua Y, Brazile BL, et al. So-called lamina cribrosa defects may mitigate IOP-induced neural tissue insult. Investigative Ophthalmology & Visual Science. 2020;61(13):15–15.

14. Zhang L, Albon J, Jones H, et al. Collagen microstructural factors influencing optic nerve head biomechanics. Investigative ophthalmology & visual science. 2015;56(3):2031–2042.

15. Brooks D, Samuelson D, Gelatt K. Ultrastructural changes in laminar optic nerve capillaries of beagles with primary open-angle glaucoma. American journal of veterinary research. 1989;50(6):929–935.

16. Fechtner RD, Weinreb RN. Mechanisms of optic nerve damage in primary open angle glaucoma. Survey of ophthalmology. 1994;39(1):23–42.

17. Quigley HA, McKinnon SJ, Zack DJ, et al. Retrograde axonal transport of BDNF in retinal ganglion cells is blocked by acute IOP elevation in rats. Investigative ophthalmology & visual science. 2000;41(11):3460–3466.

18. Carichino L, Guidoboni G, Arieli Y, Siesky BA, Harris A. Effect of lamina cribrosa deformation on the hemodynamics of the central retinal artery: a mathematical model. Investigative Ophthalmology & Visual Science. 2012;53(14):2836–2836.

19. Geijer C, Bill A. Effects of raised intraocular pressure on retinal, prelaminar, laminar, and retrolaminar optic nerve blood flow in monkeys. Investigative ophthalmology & visual science. 1979;18(10):1030–1042.

20. Beach J, Ning J, Khoobehi B. Oxygen saturation in optic nerve head structures by hyperspectral image analysis. Current Eye Research. 2007;32(2):161–170.

21. Carreau A, Hafny-Rahbi BE, Matejuk A, Grillon C, Kieda C. Why is the partial oxygen pressure of human tissues a crucial parameter? Small molecules and hypoxia. Journal of cellular and molecular medicine. 2011;15(6):1239–1253.

22. Ferrez P, Chamot S, Petrig B, Pournaras C, Riva C. Effect of visual stimulation on blood oxygenation in the optic nerve head of miniature pigs: a pilot study. Klinische Monatsblätter für Augenheilkunde. 2004;221(05):364–366.

23. Ortiz-Prado E, Dunn JF, Vasconez J, Castillo D, Viscor G. Partial pressure of oxygen in the human body: a general review. American journal of blood research. 2019;9(1):1.

24. Mari JM, Strouthidis NG, Park SC, Girard MJ. Enhancement of lamina cribrosa visibility in optical coherence tomography images using adaptive compensation. Investigative ophthalmology & visual science. 2013;54(3):2238–2247.

25. Lee EJ, Han DK, Roh YJ, Kim T-W. Underlying Microstructure of the Lamina Cribrosa at the Site of Microvasculature Dropout. Investigative Ophthalmology & Visual Science. 2024;65(8):47–47. doi:10.1167/iovs.65.8.47

26. Numa S, Akagi T, Uji A, et al. Visualization of the lamina cribrosa microvasculature in normal and glaucomatous eyes: a swept-source optical coherence tomography angiography study. Journal of Glaucoma. 2018;27(11):1032–1035.

27. Pi S, Hormel TT, Wei X, et al. Retinal capillary oximetry with visible light optical coherence tomography. Proceedings of the National Academy of Sciences. 2020;117(21):11658–11666.

28. Rao HL, Pradhan ZS, Weinreb RN, et al. A comparison of the diagnostic ability of vessel density and structural measurements of optical coherence tomography in primary open angle glaucoma. PloS one. 2017;12(3):e0173930.

29. Qian X, Kang H, Li R, et al. In vivo visualization of eye vasculature using super-resolution ultrasound microvessel imaging. IEEE Transactions on Biomedical Engineering. 2020;67(10):2870–2880.

30. Li Y, Cheng H, Duong TQ. Blood-flow magnetic resonance imaging of the retina. Neuroimage. 2008;39(4):1744–1751.

31. Causin P, Guidoboni G, Malgaroli F, Sacco R, Harris A. Blood flow mechanics and oxygen transport and delivery in the retinal microcirculation: multiscale mathematical modeling and numerical simulation. Biomechanics and modeling in mechanobiology. 2016;15(3):525–542.

32. Hua Y, Lu Y, Walker J, et al. Eye-specific 3D modeling of factors influencing oxygen concentration in the lamina cribrosa. Experimental Eye Research. 2022:109105.

33. Causin P, Guidoboni G, Harris A, Prada D, Sacco R, Terragni S. A poroelastic model for the perfusion of the lamina cribrosa in the optic nerve head. Mathematical biosciences. 2014;257:33–41.

34. Walker JA, Hua Y, McDonald H, Pallares P, Brazile B, Sigal IA. Factors influencing oxygen concentration in the lamina cribrosa. Investigative Ophthalmology & Visual Science. 2020;61(7):632–632.

35. Lee P-Y, Hua Y, Brazile BL, Yang B, Wang L, Sigal IA. A Workflow for Three-Dimensional Reconstruction and Quantification of the Monkey Optic Nerve Head Vascular Network. Journal of Biomechanical Engineering. 2022;144(6):061006.

36. Zhong F, Wang B, Wei J, et al. A high-accuracy and high-efficiency digital volume correlation method to characterize in-vivo optic nerve head biomechanics from optical coherence tomography. Acta Biomaterialia. 2022;143:72–86.

37. Tran H, Jan N-J, Hu D, et al. Formalin fixation and cryosectioning cause only minimal changes in shape or size of ocular tissues. Scientific reports. 2017;7(1):1–11.

38. An D, Pulford R, Morgan WH, Yu D-Y, Balaratnasingam C. Associations between capillary diameter, capillary density, and microaneurysms in diabetic retinopathy: A high-resolution confocal microscopy study. Translational vision science & technology. 2021;10(2):6–6.

39. Hayreh SS. Blood supply of the optic nerve head and its role in optic atrophy, glaucoma, and oedema of the optic disc. The British journal of ophthalmology. 1969;53(11):721.

40. Camasão D, Mantovani D. The mechanical characterization of blood vessels and their substitutes in the continuous quest for physiological-relevant performances. A critical review. Materials Today Bio. 2021;10:100106.

41. Shilo M, Gefen A. Identification of capillary blood pressure levels at which capillary collapse is likely in a tissue subjected to large compressive and shear deformations. Computer methods in biomechanics and biomedical engineering. 2012;15(1):59–71.

42. Lu Y, Hua Y, Wang B, et al. The Robust Lamina Cribrosa Vasculature: Perfusion and Oxygenation Under Elevated Intraocular Pressure. Investigative Ophthalmology & Visual Science. 2024;65(5):1–1. doi:10.1167/iovs.65.5.1

43. Pries AR, Secomb TW. Microvascular blood viscosity in vivo and the endothelial surface layer. American Journal of Physiology-Heart and Circulatory Physiology. 2005;289(6):H2657–H2664.

44. Pries AR, Secomb TW. Blood flow in microvascular networks. Microcirculation. Elsevier; 2008:3–36.

45. Ebrahimi S, Bagchi P. A computational study of red blood cell deformability effect on hemodynamic alteration in capillary vessel networks. Scientific reports. 2022;12(1):4304.

46. Williams J, Turney B, Moulton D, Waters S. Effects of geometry on resistance in elliptical pipe flows. Journal of Fluid Mechanics. 2020;891:A4.

47. Popel AS. Theory of oxygen transport to tissue. Critical reviews in biomedical engineering. 1989;17(3):257.

48. Lu Y, Hu D, Ying W. A fast numerical method for oxygen supply in tissue with complex blood vessel network. PloS one. 2021;16(2):e0247641.

49. Leach R, Treacher D. Oxygen transport2. Tissue hypoxia. Bmj. 1998;317(7169):1370-1373.

50. Loiacono LA, Shapiro DS. Detection of hypoxia at the cellular level. Critical care clinics. 2010;26(2):409–421.

51. McKeown S. Defining normoxia, physoxia and hypoxia in tumours—implications for treatment response. The British journal of radiology. 2014;87(1035):20130676.

52. Selbach MJ, Wonka F, Hoper J, Funk RH. Effects of elevated intraocular pressure on haemoglobin oxygenation in the rabbit optic nerve head: a microendoscopical study. Exp Eye Res. Sep 1999;69(3):301–9. doi:10.1006/exer.1999.0702

53. Hockel M, Vaupel P. Tumor hypoxia: definitions and current clinical, biologic, and molecular aspects. Journal of the National Cancer Institute. 2001;93(4):266–276.

54. Ortiz-Prado E, Natah S, Srinivasan S, Dunn JF. A method for measuring brain partial pressure of oxygen in unanesthetized unrestrained subjects: the effect of acute and chronic hypoxia on brain tissue PO2. Journal of neuroscience methods. 2010;193(2):217–225.

55. Secomb TW, Hsu R, Park EY, Dewhirst MW. Green’s function methods for analysis of oxygen delivery to tissue by microvascular networks. Annals of biomedical engineering. 2004;32(11):1519–1529.

56. Harris A, Ciulla TA, Chung HS, Martin B. Regulation of retinal and optic nerve blood flow. Archives of ophthalmology. 1998;116(11):1491–1495.

57. Wang L, Burgoyne CF, Cull G, Thompson S, Fortune B. Static blood flow autoregulation in the optic nerve head in normal and experimental glaucoma. Investigative Ophthalmology & Visual Science. 2014;55(2):873–880.

58. Sun X, Dai Y, Chen Y, et al. Primary angle closure glaucoma: what we know and what we don’t know. Progress in retinal and eye research. 2017;57:26–45.

59. Xuan M, Wang W, Bulloch G, et al. Impact of Acute Ocular Hypertension on Retinal Ganglion Cell Loss in Mice. Translational Vision Science & Technology. 2024;13(3):17–17. doi:10.1167/tvst.13.3.17

60. Tun TA, Thakku SG, Png O, et al. Shape Changes of the Anterior Lamina Cribrosa in Normal, Ocular Hypertensive, and Glaucomatous Eyes Following Acute Intraocular Pressure Elevation. Investigative Ophthalmology & Visual Science. 2016;57(11):4869–4877. doi:10.1167/iovs.16-19753

61. Yu J, Liang Y, Thompson S, Cull G, Wang L. Parametric transfer function analysis and modeling of blood flow autoregulation in the optic nerve head. International Journal of Physiology, Pathophysiology and Pharmacology. 2014;6(1):13.

62. Hayreh S. The optic nerve head circulation in health and disease. Ophthalmic Literature. 1996;2(49):111.

63. Hayreh SS. Blood flow in the optic nerve head and factors that may influence it. Progress in retinal and eye research. 2001;20(5):595–624.

64. Mackenzie PJ, Cioffi GA. Vascular anatomy of the optic nerve head. Canadian Journal of Ophthalmology. 2008;43(3):308–312.

65. Mozaffarieh M, Grieshaber MC, Flammer J. Oxygen and blood flow: players in the pathogenesis of glaucoma. Molecular vision. 2008;14:224.

66. Orgül S, Gugleta K, Flammer J. Physiology of perfusion as it relates to the optic nerve head. Survey of ophthalmology. 1999;43:S17–S26.

67. Waxman S, Brazile BL, Yang B, et al. Lamina cribrosa vessel and collagen beam networks are distinct. Experimental Eye Research. 2022;215:108916.

